# Characterization of the direct and indirect inhibition of apoptosis by full-length recombinant Bcl-xL monomers

**DOI:** 10.1101/2025.06.16.659449

**Authors:** Christina Elsner, Ludovica M. Epasto, Adeline Cieren, Dominik Gendreizig, Svetlana Kucher, Daniel Roderer, Enrica Bordignon

**Affiliations:** Department of Physical Chemistry, Sciences II, University of Geneva, 30 Quai Ernest Ansermet, 1211 Geneva, Switzerland; Leibniz-Forschungsinstitut für Molekulare Pharmakologie, Robert-Rössle-Str. 10, 13125 Berlin, Germany

## Abstract

The Bcl-2 protein Bcl-xL is an inhibitor of intrinsic apoptosis which either directly inhibits the pore-forming Bcl-2 proteins, like Bax or Bak, or indirectly inhibits pore formation by sequestering the pro-apoptotic BH3-only activators. The structural basis of the inhibition of pore formation in the outer mitochondrial membrane is still largely unknown due to the lack of atomic resolution structures of the relevant inhibitory complexes at the membrane. Here we present a protocol to obtain high-yield recombinant monomeric full-length Bcl-xL proteins. The monomeric Bcl-xL retains the ability to shuttle between membrane and aqueous environments and can successfully inhibit Bcl-2-induced membrane permeabilization via both modes of action, as proven by *in vitro* and *in organelle* assays with a minimal Bcl-2 interactome constituted by Bcl-xL, cBid, and Bax.

## Introduction

Regulation of apoptosis plays a major role in cellular homeostasis and the immune system and any interference in the activation and inhibition mechanisms gives rise to diseases like cancer and Alzheimer’s [1]. In the mitochondrial pathway of apoptosis such regulation is finely tuned by the members of the Bcl-2 protein family. The effector or pore-forming proteins (Bax, Bak, Bok) cause mitochondrial outer membrane permeabilization (MOMP) triggering a signal cascade ultimately leading to apoptosis. This process is activated by BH3-only proteins (called also activators, such as cBid, Bim, PUMA, or sensitizers, such as Bad, Noxa, etc.) and inhibited by anti-apoptotic proteins (called also pro-survival or guardians, such as Bcl-xL, Bcl-2, etc.). The complex interplay between these three sub-groups of the Bcl-2 protein family, which will ultimately decide the fate of the cell, mostly requires the presence of the mitochondrial outer membrane as active player (for recent reviews see [2–4]). The Bcl-2 interactome is regulated by diverse interactions between Bcl-2 homology (BH) domains and most proteins also contain a hydrophobic, putative transmembrane (TM) C-terminal region. As their name implies, BH3-only proteins only have the BH3-domain and a TM C-terminal helix (TMH) and are mostly intrinsically disordered, with the notable exception of Bid, which has a globular fold and lacks the C-terminal TMH[5]. Both pore formers and pro-survival proteins have four conserved BH domains (1-4) and the C-terminal TMH, the latter being often cleaved off in recombinant proteins (ΔC protein variants) to increase their stability in *in vitro* studies. However, the C-terminal hydrophobic sequences were shown to have not only the ability to target the proteins to the membrane, but also to engage in protein-protein regulatory interactions[4]. Interactions between Bcl-2 proteins are often mediated by the conserved hydrophobic groove assembled by the BH-domains 1-3 of pore formers and pro-survival proteins, which can efficiently bind BH3-motifs of other Bcl-2 proteins[2]. Besides the BH3-into-groove interaction, BH3-domains of proapoptotic proteins were found to bind the rear side of Bax (*α*1/*α*6 helices)[6] inducing its activation. Additionally, full-length Bax and Bcl-xL were found to interact via *α*1-helices [7], or via three-dimensional domain swapping [8, 9].

The interaction between pro-survival and pro-apoptotic proteins is the molecular basis for the inhibition of MOMP. Three different inhibition modes have been described in literature: the guardians can regulate the localization of the effectors to reduce binding to the MOM, they can bind the pore-formers at the membrane (direct mode), or they can sequester the BH3-only activators (indirect mode) [3, 6, 10, 11]. The latter mode is especially interesting, since it has been shown that Bcl-xL has a higher affinity to BH3-only proteins than to pore formers [2].

The study of these inhibitory complexes is complicated by the challenges in preparing active, full-length recombinant proteins at high concentrations and which can shuttle between their soluble and membrane-bound states[21, 22]. Experimental atomic resolution structures of Bcl-xL exist, but only of various deletion-variants. In particular, the structure of Bcl-xL in water was solved by NMR on a ΔC variant (PDB entry 1LXL, (see fig 1A)), the BH3-to-groove binding was characterized on deletion-variants of the long loop region of Bcl-xL (e.g. Δ45-84) and of the C-terminal transmembrane domain (e.g. Δ212-233), in complex with peptides derived from BH3-domains (see e.g. PDB entries: 4QVE, 1BXL, 1G5J, 3FDL, 4CIN). Biophysical studies of full-length Bcl-xL in its soluble or membrane-anchored form have shed light on the role of the C-terminal TMH in anchoring Bcl-xL to the membrane and binding some BH3-only proteins (e.g. Bim), and have validated the ability of membrane-bound Bcl-xL to bind BH3 domains with its hydrophobic groove. Table 1 summarizes the available purification protocols for full-length Bcl-xL and the methods used to study their properties in solution and/or in membranes.

**Table 1:** Examples of available protocols for the preparation of full-length Bcl-xL (no PDB is available for full-length Bcl-xL).

| Sequence length | Key purification details | Membrane interaction | Soluble protein | Activity assay | Methods and Ref |
| --- | --- | --- | --- | --- | --- |
| GEF-1-233 | 1. His-GB1-Tag; 2. SEC | LUVs, pH-induced membrane insertion | yes | no | CD, fluorescence [12] |
| 1-233 | 1. C-terminal intein-chitin binding domain; 2. Anion-exchange chromatography | LUVs | yes | yes | Fluorescence [13, 14] |
| His-Tag-1-233 | 1. His-Tag; 2. SEC | no | yes | no | Purification [15], Fluorescence, EPR [16] |
| 1-233 | 1. C-terminal intein-chitin binding domain; 2. Phenyl Sepharose column | LUVs and mitochondria | yes | yes | Fluorescence, e.g. [17, 18] |
| 1-199-LPETG-202-233 | 1.TM helix: His-Tag, precipitation, solubilization. 2.Soluble domain: His-Tag, SEC | reconstitution of TMH and ligation of the soluble domain | no | no | NMR [19] |
| 1-233 | 1. C-terminal intein-chitin binding domain; 2. Minor fraction soluble, insoluble fraction treated with urea and SDS; 3. SEC | reconstitution into nanodiscs | yes | no | NMR and MD simulations [20] |
| 1-233 | 1. C-terminal intein-chitin binding domain; 2. Anion exchange chromatography 3. SEC | LUVs and mitochondria | yes | yes | EPR, this work |

Examples exist of monomeric full-length Bcl-xL in solution and anchored to nanodiscs (fig 1A [20]), however truncated and full-length Bcl-xL have also been described to form dimers and higher order oligomers (see some examples in fig 1A), which were previously discussed to play a potential role in apoptosis [8, 14, 18, 23, 24].

In this work we present a simple purification protocol which allows efficient separation of full-length Bcl-xL monomers (residues 1-233) from the oligomeric fraction which is formed during purification (in the following named Bcl-xL^O^). The obtained recombinant monomeric Bcl-xL is stable and fully active, retaining the ability to shuttle between membrane and solution and to perform direct and indirect inhibition of MOMP, as demonstrated by electron paramagnetic resonance (EPR) spectroscopy and various activity assays performed on a minimal Bcl-2 interactome *in vitro* and *in organelle*.

## Results

### Characterization of Bcl-xL: monomers and oligomers

In the presented protocol we used a Bcl-xL construct containing a cleavable C-terminal intein tag with a chitin binding domain (Table 1), which was also successfully used in several other publications to purify full-length Bcl-xL (see for example [17, 18, 20]). After affinity chromatography and tag cleavage, anion-exchange chromatography was performed (Table 1) which was found to be necessary to achieve a high activity of the final monomeric fraction. A final size exclusion chromatography step was added to isolate the monomeric fraction from the oligomeric fraction (fig 1B, see also materials & methods and fig. S1, S2, S3B). The oligomeric fraction was further investigated by mass photometry and it was found to be heterogeneous in size with an average molecular weight of about 250 kDa (fig 1C). At nanomolar concentrations, the oligomers were found to dissociate into smaller particles, likely monomers and/or dimers (fig. 1C). Negative stain electron microscopy (NS-EM) performed on the isolated Bcl-xL^O^ confirmed the dissociation into smaller units (see fig. S4). The dissociation of the oligomers at nanomolar concentrations, and the fact that the oligomeric fraction is observed only after the anion exchange column (fig. S1 and S2, also previously observed in [8]), favor the hypothesis that it only has a minor or no physiological role.

**Figure 1:**
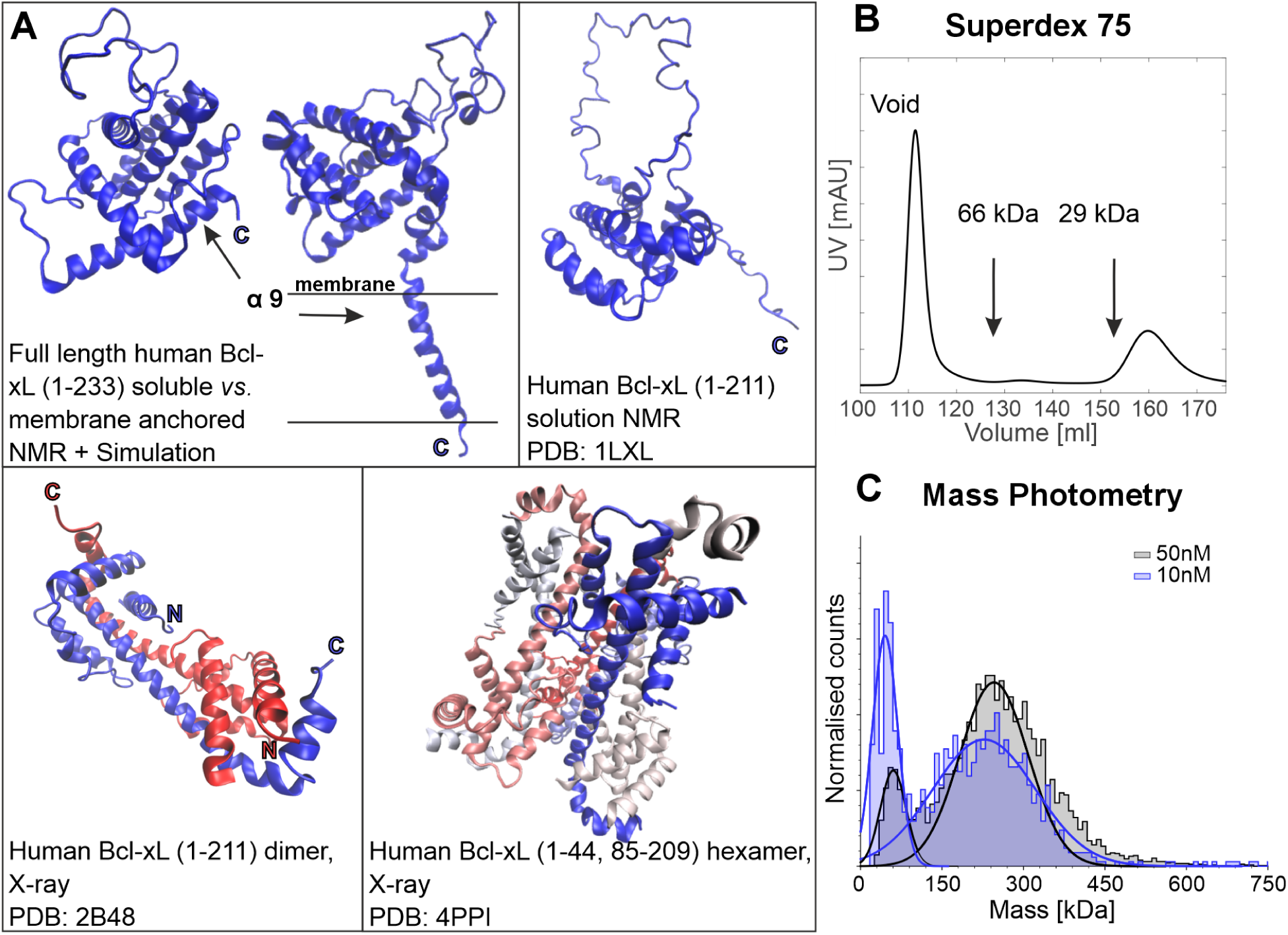
Monomeric, dimeric, and oligomeric forms of Bcl-xL. **A:** Examples of Bcl-xL structures. Upper panel left: full-length Bcl-xL in aqueous and nanodisc environment obtained by NMR and MD simulations (atomic coordinates kindly provided by Prof. Marassi from [20]); right: NMR structure of monomeric Bcl-xLΔC in solution; lower panel left: crystal structure of a water-soluble dimer of Bcl-xLΔC formed by 3D-domain swapping; right: crystal structure of a water-soluble hexamer of Bcl-xLΔC-Δloop formed by domain-swapped dimers. **B:** Size exclusion profile from our Bcl-xL purification protocol performed on a Superdex 75 26/600 column at 4°C. Arrows indicate reference protein elution volumes. The peak eluting in the void volume corresponds to Bcl-xL^O^ (size *>*70 kDa), the peak eluting at 160 ml corresponds to monomeric Bcl-xL (molecular weight of 26 kDa). Additional SEC analysis can be found in fig. S1 **C:** Mass photometry of Bcl-xL^O^ at 10 nM (blue) and 50 nM (grey) (protomer concentrations). For the 10 nM sample, we found a low-molecular-mass peak close to the mass sensitivity limit of the method (30 kDa) centered around 45 kDa (*σ*=23 kDa, 34% fraction) and a high-molecular-mass peak at 228 kDa (*σ*=95 kDa, 77% fraction). For the 50 nM sample, the low-molecular-mass peak is centered at 60 kDa (*σ*=24 kDa, 14% fraction) and the high-molecular-mass peak at 240 kDa (*σ*=82 kDa, 81% fraction).

**Figure 2:**
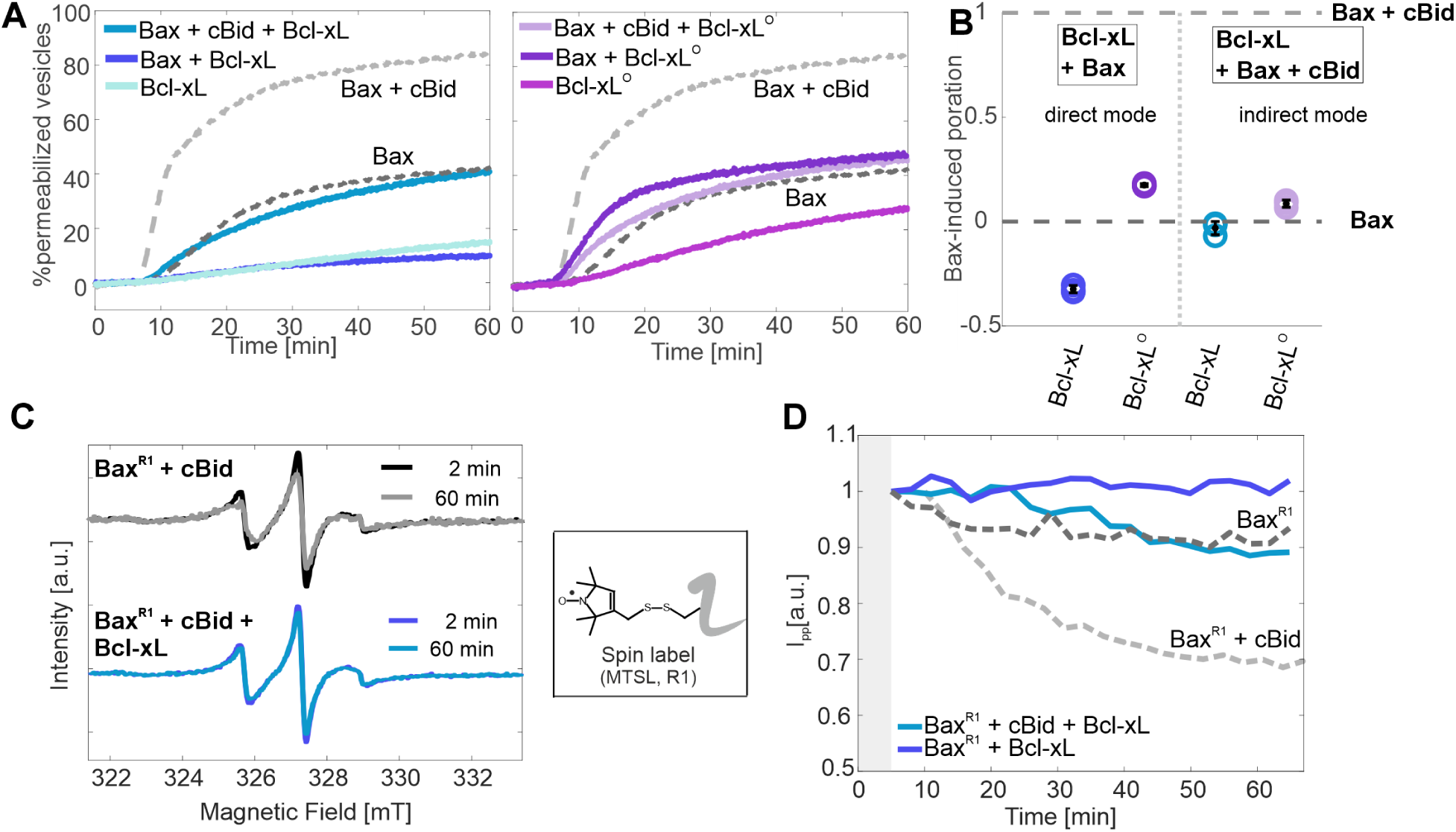
*In vitro* fluorescence and EPR activity assays. **A:** Examples of pore formation assays based on the release of the fluorescent probe calcein from LUVs through Bax-induced pores. The time traces show the amount of permeabilized vesicles at 37°C over time, normalized to the highest permeabilization induced by lysis with detergents (100%) (see materials & methods and fig. S3). All proteins were added at a concentration of 25 nM. The dotted grey lines on the two graphs show reference measurements (Bax alone, dark grey, 40% autoactivity; Bax + cBid, light grey, 80% activity). Left: effects induced by monomeric Bcl-xL. Right: effects induced by Bcl-xL^O^. **B:** Comparison of 3 technical replicates of the inhibitory effects shown in A. Data are taken after 30 min of incubation, with 1 being the normalized maximal Bax activity in presence of cBid and 0 Bax autoactivity for each replicate. **C:** EPR activity assay on the monomeric Bcl-xL. Examples of the changes in EPR spectra of Bax^R1^ in absence (top) and presence of monomeric Bcl-xL (bottom) at different incubation times at 37°C with large unilamellar vesicles (LUVs) and cBid. The inset shows the spin label MTSL attached to a cysteine yielding the residue R1. The decrease of the spectral intensity reflects Bax oligomerization and membrane insertion, which is inhibited by Bcl-xL. **D:** EPR kinetics of the membrane insertion of Bax^R1^ obtained by plotting the intensity of the central line of the EPR spectra (I_pp_) over time. Both the cBid-induced Bax activity (light grey, dashed) and the Bax autoactivity (dark grey, dashed) are inhibited by monomeric Bcl-xL (light and dark blue, respectively).

### Direct and indirect inhibition of apoptosis ***in vitro*** and ***in organelle***

To further shed light on the physiological role of the two isolated fractions of Bcl-xL, we used a minimal Bcl-2 interactome consisting of cBid, Bax, and either monomeric Bcl-xL or Bcl-xL^O^. With near physiological protein concentrations (25 nM) [2], we tested Bcl-xL’s ability to perform direct and indirect inhibition of Bax-induced pore formation *in vitro* by monitoring the leakage of highly concentrated, self-quenching calcein from LUVs (fig. 2A,B, see also materials & methods and fig. S3). To be able to detect both modes of inhibition on the same batch of LUVs, we optimized the Bax-to-lipid ratio to reach a high level of autoactivity (about 40%, dark grey dashed lines in fig. 2A) while maintaining a good contrast to the maximal cBid-induced Bax activity (Bax:cBid 1:1 stoichiometric ratio) (80% permeabilization, light grey dashed lines, in fig. 2A). Under these conditions, control experiments with Bcl-xL monomers showed a 15% background calcein leakage, while Bcl-xL^O^ induced a higher leakage of about 30% after 60 min. We proved the direct inhibition adding either monomeric or oligomeric Bcl-xL at stoichiometric ratio in presence of Bax (1:1 protein ratio) to test the inhibition of Bax autoactivity. Monomeric Bcl-xL efficiently inhibited Bax autoactivity to values of around 10% (blue in fig. 2A). In contrast, Bcl-xL^O^ failed to inhibit Bax autoactivity (dark purple in Fig. 2A). Figure 2B summarizes the results on three technical repeats. By adding either monomeric or oligomeric Bcl-xL at stoichiometric ratio in presence of Bax and cBid (1:1:1 protein ratio) we found that calcein release was decreased to a value similar to Bax autoactivity (around 40%) (cyan and light purple in fig. 2A), indicating that both Bcl-xL monomers and the heterogeneous mixture of Bcl-xL^O^ can efficiently inhibit cBid at stoichiometric ratio (for the oligomers, we consider the protomer stoichiometry) at nanomolar concentration. However, due to the heterogeneity of Bcl-xL^O^ (see fig. 1C and fig. S4), it is not possible to disentangle the role of the higher order oligomers from that of the smaller units present in solution. The *in vitro* activity tests demonstrate that the purified monomeric full-length Bcl-xL retains the ability to efficiently inhibit cBid as well as Bax at molar stoichiometric ratios, while Bcl-xL^O^ retains only the indirect mode of action.

We corroborated the activity data of the monomeric Bcl-xL also at micromolar concentrations via EPR kinetics experiments following the membrane insertion of MTSL-labeled wild type Bax (called for simplicity Bax^R1^) (fig. 2C,D) in presence and absence of monomeric unlabeled Bcl-xL (EPR method previously shown in [25, 26]). This method allows monitoring changes in Bax dynamics in contrast to following leakage from the LUVs. Notably, Bcl-xL^O^ could not be tested via EPR kinetics due to the too low concentration in the stock solution. The decrease of the EPR spectral intensity (central line in fig. 2C) plotted over time (fig. 2D) indicates the reduced mobility of spin-labeled Bax which can arise from it entering the membrane and oligomerizing either alone (autoactivity, dotted dark grey) or in presence of cBid (dotted light grey). If Bcl-xL monomers are added at stoichiometric ratio, they inhibit Bax autoactivity (dark blue) and cBid (light blue), fully in agreement with the fluorescence data in fig. 2A,B. The fluorescence and EPR *in vitro* assays prove that purified monomeric full-length Bcl-xL inhibits Bax and cBid both at nano- and micromolar concentrations.

To confirm the physiological relevance of the direct and indirect modes of action of the purified monomeric Bcl-xL, we performed activity tests using freshly isolated mitochondria from mouse liver (see fig. 3A and materials & methods). We monitored the release of cytochrome c upon incubation with Bax in presence of cBid and/or Bcl-xL. The maximum absorption intensity of the cytochrome c released in the supernatant was used as an indicator of MOMP. Figure 3B shows examples of the UV-vis spectra obtained under different conditions (see materials & methods) and 3C the statistical analysis over several biological replicates. We found that mitochondria alone have a detectable release of cytochrome c, which we define as background release (3B and dashed dark grey line at 0 in 3C) and that the maximal release is obtained in presence of Bax and cBid at 1:0.1 stoichiometric ratio (3B and dashed light grey line at 1 in 3C). In contrast to the *in vitro* assays, cBid was not used 1:1 with Bax to minimize effects of its own activity and its interaction with endogenous Bax/Bak. Bcl-xL alone does not induce MOMP above the background level observed (cyan circles in 3C), but cBid alone has a clear effect in membrane permeabilization (magenta circles in 3C). The effect observed with cBid can be either due to its own ability to permeabilize membranes [27] and/or to the activation of the endogenous Bax/Bak present in the mitochondria. Interestingly, when Bcl-xL is added together with cBid, the cytochrome c release goes back to background levels (violet circles in 3C), indicating that Bcl-xL efficiently inhibits cBid-induced MOMP. Bax alone (green circles in 3C) also induces cytochrome c release, and this effect can be fully inhibited by addition of Bcl-xL already in a 1:0.1 Bax:Bcl-xL stoichiometric ratio (compare orange and brown circles in 3C). To fully inhibit the maximal permeabilization with Bax and cBid, it was not sufficient to add Bcl-xL at stoichiometric ratio to cBid (1:0.1:0.1, light blue circles in 3C). However, a higher Bcl-xL concentration, equal to the sum concentration of Bax and cBid, resulted in full inhibition (1:0.1:1.1 ratio, blue circles in 3C). The activity assay performed in mitochondria corroborated the conclusions obtained *in vitro* and validated the physiological relevance of the monomeric full-length Bcl-xL, paving the way towards its use for biophysical analysis.

**Figure 3:**
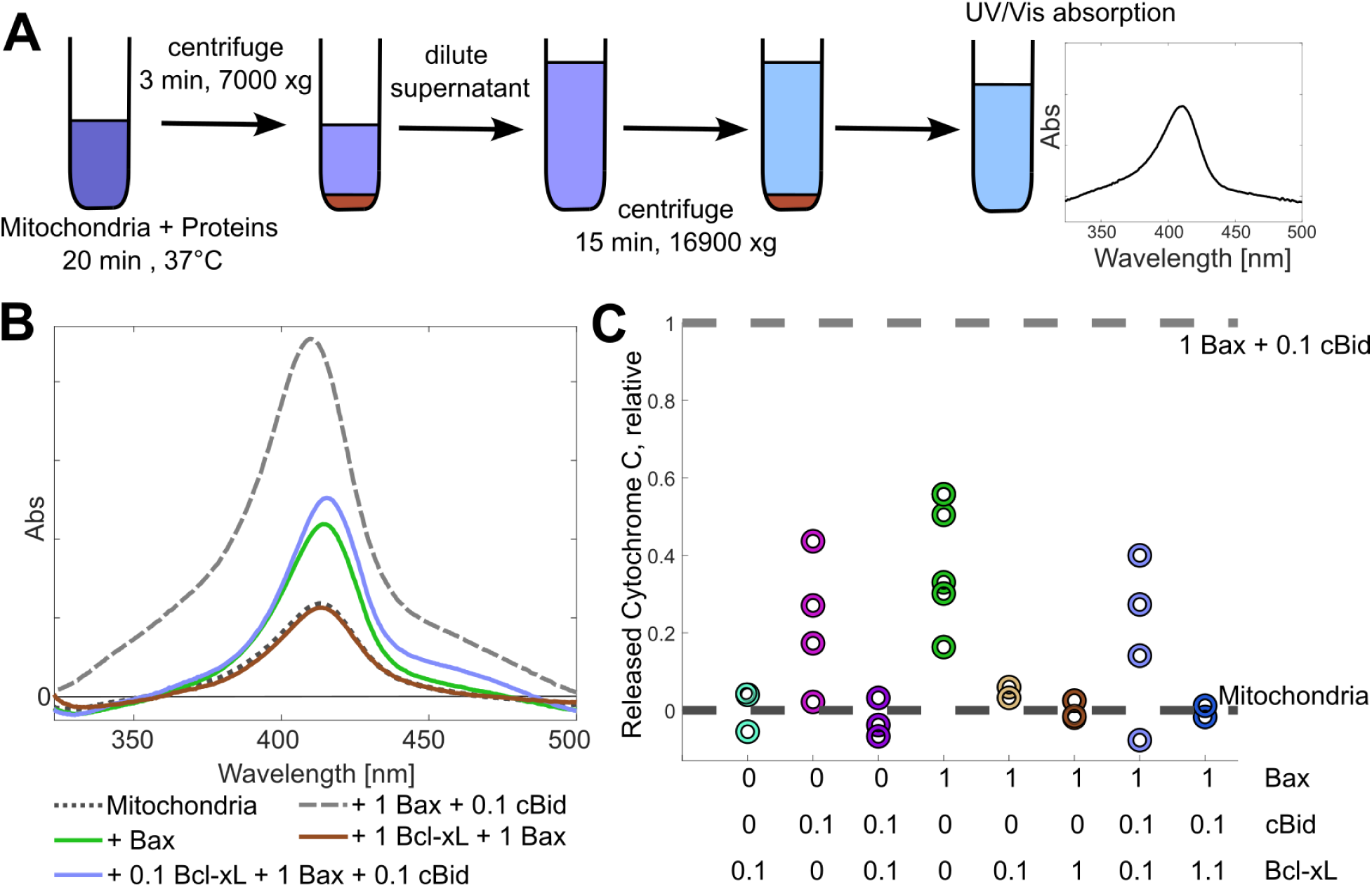
*In organelle* activity assays of monomeric Bcl-xL. **A:** Schematic description of cytochrome c release assays by UV-vis spectroscopy (see materials & methods). Mitochondria and proteins are incubated with freshly isolated mitochondria and the supernatant is then separated from the pellet via centrifugation steps. Membrane permeabilization is monitored by the intensity of the UV-vis absorption of the released cytochrome c (408-414 nm for the oxidized and reduced form) **B:** Example of the background corrected spectra obtained with one mitochondria batch under different conditions. **C:** Statistical analysis of the cytochrome c release in presence of different combinations of proteins (biological replicates). The value 1 corresponds to the maximum release caused by incubation with Bax:cBid (1:0.1 ratio) on the mitochondria batch under consideration, the value 0 is the release caused by incubating the respective mitochondria batch alone.

### Water-membrane partitioning of Bcl-xL monomers and interactions with Bcl-2 partners

To understand the role of the membrane in the formation of the inhibitory complexes, we characterized the partitioning of Bcl-xL between aqueous solution and membrane environment using SDS PAGE and EPR spectroscopy. In addition, we studied the effect exerted on the water-membrane equilibrium of Bcl-xL by other Bcl-2 partners, namely cBid and Bax. To quantify the amount of Bcl-xL in both environments, we used a simple centrifugation method: we incubated Bcl-xL and LUVs in the presence or absence of cBid or Bax for one hour at 37°C, followed by separation of the mixture via centrifugation (fig. 4A). The pellet (P) and supernatant (S) fractions were separately analyzed (fig. 4B,C).

**Figure 4:**
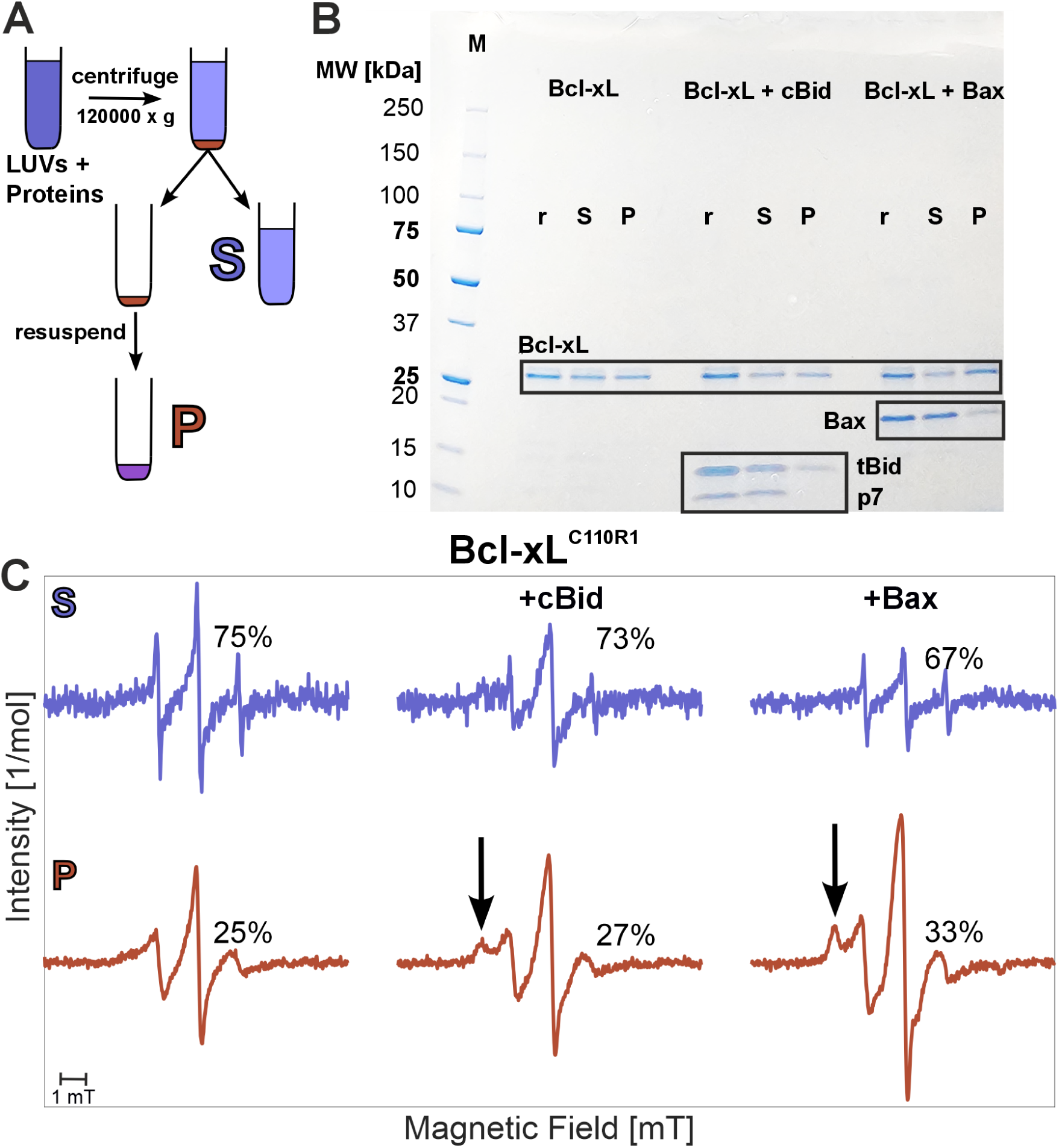
Membrane affinity of monomeric Bcl-xL in absence and presence of partner Bcl-2 proteins. **A:** Schematic description of the sample preparation. The protein-LUV mixture is incubated for one hour at 37°C. The pellet and supernatant fractions are separated by centrifugation (see materials & methods). **B:** SDS PAGE analysis of Bcl-xL alone or in presence of cBid or Bax and LUVs. *P*: pellet; *S*: supernatant; *r*: reference protein mixture in solution without LUVs. Boxes highlight bands corresponding to monomeric Bcl-xL (26 kDa), Bax (21 kDa), cBid (tBid and p7 bands at 15 and 7 kDa, respectively).**C:** EPR spectra of the supernatant and pellet fractions obtained from mixtures of LUVs with Bcl-xL^C110R1^ alone (left) or in combination with cBid (middle) or Bax (right) in a 1:1 stoichiometric ratio at micromolar concentration. The percentages indicate the relative amount of moles (10% estimated error) of spin-labeled Bcl-xL in each fraction obtained by spectral integration. The arrows point to spectral changes caused by the interaction of Bcl-xL with Bcl-2 partners in the membrane.

The SDS PAGE analysis shown in fig. 4B confirms the equilibrium of Bcl-xL between its membrane-bound and soluble states, corroborating previous data *in vivo*, showing that about 50% Bcl-xL is cytoplasmic[21]. The presence of cBid does not affect this equilibrium. Notably, in the supernatant both bands of the cleaved cBid (tBid, 15 kDa and p7, 7 kDa) can be detected, however, the pellet fraction shows only tBid. If full-length Bax is present, the amount of Bcl-xL at the membrane slightly increases and most of Bax is found in the supernatant.

To investigate Bcl-xL in solution and in membrane by EPR, we engineered an accessible cysteine at position 110 and spin labeled it with MTSL (Bcl-xL^C110R1^). After labeling, the proteins retain their activity, as proven by pore formation assays (see fig. S3). The label is strategically positioned in the hydrophobic groove of Bcl-xL, which was previously shown to bind BH3-peptides[8, 23]. The EPR spectra of Bcl-xL^C110R1^ recorded in the supernatant and pellet fractions allowed quantification of the labeled proteins using the double integral of the spectra, proportional to the concentration of Bcl-xL^C110R1^ (fig. 4C). We found that 25% of the total amount of Bcl-xL monomers binds to the membrane. This partitioning remains unchanged in the presence of cBid. In the presence of Bax the amount of membrane-bound Bcl-xL slightly increased to 33%, in line with the result from SDS PAGE (fig. 4B). Notably, the spectral shape of Bcl-xL in the membrane changed when cBid or Bax were present during incubation. The arrows in fig. 4C highlight the broad spectral component emerging in the presence of the Bcl-2 partners, which indicates that the motion of the spin-labeled side chain in the hydrophobic groove of Bcl-xL^C110R1^ is slowed down due to sterical hindrance, possibly caused by a direct interaction with the other Bcl-2 partners (formation of hetero-complexes) and/or by structural changes within Bcl-xL triggered by the presence of the interaction partner.

## Discussion

The presented optimized purification protocol of recombinant full-length Bcl-xL yielded a pure, fully active water-soluble monomeric Bcl-xL well separated from a fraction of higher order oligomers. The oligomers, which are formed during the purification procedure, easily aggregate upon concentration, partially dissociate into smaller units at nanomolar concentrations and can execute only the indirect mode of inhibition. The propensity of truncated variants of Bcl-xL to form dimers and higher-order oligomers and the difficulty in avoiding large oligomers when purifying full-length Bcl-xL prompted speculations about their possible role in the inhibition of apoptosis. Notably, it was shown for the 3D domain-swapped dimer formed by truncated Bcl-xL variants, that the BH3-binding groove remains accessible to BH3 peptides[23], which might be sufficient to retain the ability to indirectly inhibit pore formation via sequestration of BH3-only proteins. Here we found that the oligomers can also target the membrane (fig. S5), which will possibly facilitate the interaction with tBid, explaining the observed indirect inhibition pathway.

In contrast, the isolated monomeric fraction is stable, and can be concentrated up to 10-30 *µ*M, which is favorable for further biophysical studies. Monomeric Bcl-xL is proven to execute both direct and indirect modes of inhibition, as shown by assays using a minimal interactome consisting of Bcl-xL, cBid, and Bax. We used established *in vitro* pore formation assays, which provides standardized activity tests of protein batches and complemented them with *in organelle* assays that have a higher physiological relevance, still allowing fast detection of the activity of different combinations of Bcl-2 proteins.

With this protocol, we obtained recombinant fully active proteins at micromolar concentrations, which we could spin label without loss of activity. This paves the way towards the use of site-directed spin labeling EPR and other biophysical methods to study dynamics and structures of inhibitory complexes at the membrane. Indeed, we identified a dynamic change in the hydrophobic groove of membrane-embedded spin-labeled Bcl-xL not only in presence of tBid, but also in the presence of Bax, which is promising for the further elucidation of the structural basis of the regulatory mechanisms of Bcl-2 proteins in the mitochondrial pathway of apoptosis.

## Methods

### Expression of recombinant proteins

The plasmids pTYB1-Bcl-xLC110 (S110C-C151S) and pET23d-BidC82 (C30S-S82C-C126S) were cloned by site-specific mutagenesis and controlled via DNA sequencing. His-tagged mouse Bid (pET23d[28]), human Bax (pTYB1[29]), and human Bcl-xL (pTYB1[13]), kindly provided by Prof. Garcia-Saez were expressed in *E. coli* and induced with IPTG for 4h (Bid) or overnight (Bcl-xL, Bax) and the pellets were either used immediately or shock frozen in liquid nitrogen and stored at −80°C. After lysis of the bacteria in a microfluidizer and incubation with 1-200U DNase I per liter of bacterial culture, the membranes and intact bacteria were separated from the soluble fraction via centrifugation at 35000xg for 35 min.

### Purification and cleavage of Bid

Bid was purified from the soluble fraction of the bacterial lysate using Ni-NTA (Ni-NTA Superflow by Qiagen). It was washed first with Nickel buffer (20 mM Tris, pH 7.5, 20 mM NaH2PO4, 300 mM NaCl) without imidazole, then with 10 mM and 25 mM before being eluted with 250 mM imidazole. The protein was stored in storage buffer (20 mM Tris, pH 7.5, 150 mM NaCl).

Bid (100-150 µM) was mixed 1:1 with cleavage buffer (50 mM HEPES, pH 7.5, 100 mM NaCl, 10 mM DTT, 1 mM EDTA, 10% sucrose) and Caspase 8 (final concentration 1 µM). The mixture was incubated for 3 h at room temperature and purified using Ni-NTA, washed with Nickel buffer without imidazole, eluted with 250 mM imidazole and dialyzed back into storage buffer. In the following, the cleaved Bid is called cBid, the 15 kDa fragment of cBid is called tBid (also known as p15 in literature) and the 7 kDa fragment is called p7.

### Purification and spin labeling of Bcl-xL and Bax

Bcl-xL and Bax were purified from the soluble fraction of the bacterial lysate using a chitin resin (New England Biolabs) which was washed 20x with chitin buffer (20 mM Tris (Bcl-xL) or 8 mM Tris (Bax), pH 8.0, 500 mM NaCl) and 3x cleavage buffer (chitin buffer + 30 mM DTT freshly added). The column was filled with cleavage buffer, closed and incubated at 4°C overnight to cleave off the intein tag. The proteins were eluted with chitin buffer and dialyzed 1:10 against 20 mM Tris (pH 8.0) with one step (Bcl-xL) or 8 mM Tris (pH 8.0) in five steps (Bax), respectively.

For Bcl-xL, 1 mM DTT was added to the dialysate before loading on an anion exchange column (POROS™ GoPure™ 50 HQ Pre-packed Column) at 0.5 ml/min. The column was washed with Mono A50 buffer (20 mM Tris, pH 8.0, 50 mM NaCl, 1 mM DTT) and the protein was eluted running a gradient with Mono B buffer (20 mM Tris, pH 8.0, 1 M NaCl, 1 mM DTT). The pooled fractions of the main peak were injected into a size exclusion column (Superdex 200 10/300) equilibrated with storage buffer. If the protein had to be labeled afterwards, 10 µM TCEP was freshly added to the storage buffer. The resulting oligomer peak was pooled but not concentrated as centrifugal filters cause the oligomers to aggregate. The monomer peak was pooled and concentrated in centrifugal filters (Vivaspin® Turbo 4 Sartorius (cutoff 10 kDa)) (see fig. S1). During the optimization procedure, we found that skipping the anion exchange column and performing directly the SEC resulted in significantly higher amount of monomers with respect to oligomers; however the resulting monomers were less active (see fig. S2, S3).

Starting from a culture of 1.5 liters, we collected 4 mL of the monomeric fraction at a concentration of around 1-2 *µ*M, which was found to be stable in buffers at pH 7.5 and could be further concentrated in centrifugal units to 10-30 *µ*M with negligible protein losses. The Bcl-xL^O^ fraction (10 mL, 5-10 *µ*M protomer concentration) could not be further concentrated due to severe protein aggregation.

To label the monomeric fraction of Bcl-xL, 5-10x excess MTSL was added per protein (protein concentration 10-20 µM) and incubated overnight at 8°C. The excess label was removed by washing 4-5x in centrifugal filters (Vivaspin® Turbo 4 Sartorius (cutoff 10 kDa)) with storage buffer. The spin labeling efficiency (spin per cysteine) of monomeric Bcl-xL^C110R1^ was 49%, as calculated by comparing the second integral of the cw EPR spectra with a reference TEMPOL standard (see fig. S6).

For Bax, the dialysate was loaded on an anion exchange column (HiTrap® Q High Performance) at 0.5 ml/min. The column was washed with Mono A buffer (8 mM Tris, pH 8.0) and the protein was eluted running a gradient with Mono B buffer (20 mM Tris, pH 8.0, 1 M NaCl) pooling the first peak.

To label wild type Bax (containing two natural cysteines C62 and C126), 10x excess MTSL per protein (protein concentration 10-20 µM) was added and incubated overnight at 4°C. The excess label was removed by washing 4-5x in centrifugal filters (Vivaspin® Turbo 4 Sartorius (cutoff 10 kDa)) with storage buffer. The spin labeling efficiency of Bax^R1^ was 94% (spin per cysteine), as calculated by comparing the second integral of the cw EPR spectra with a reference TEMPOL standard (see fig. S6).

For both, Bax and Bcl-xL, we found that using the Vivaspin® Turbo 4 Sartorius centrifugal filters caused less protein loss compared to other centrifugal filters. For Bcl-xL, removing excess label in spin desalting columns resulted in complete protein loss. All proteins were tested in pore formation assays before further experimental use.

### Mass photometry

Mass photometry was measured at different concentrations on a TwoMP Mass Photometer from Refeyn (mass range 30 kDa – 5 MDa) in the Protein Production and Structure Core Facility at EPFL, Lausanne, Switzerland.

### LUVs Preparation

The lipid mixture mimicking the mitochondrial outer membrane was composed of 46% egg L-*α*-phosphatidyl choline (PC), 25% egg L-*α* phosphatidyl ethanolamine (PE), 11% bovine liver L-*α*-phosphatidyl inositol (PI), 10% 18:1 phosphatidyl serine (PS) and 8% cardiolipin (CL) (% w/w). The lipid mixture in chloroform was aliquoted, dried under vacuum, flushed with nitrogen and stored at −80°C for a maximum time of one year. To prepare large unilamellar vesicles the mix was solved in either storage buffer to give a stock solution of 20 mg/ml (for EPR experiments) or in calcein (80 mM, adjusted to pH 7.0 with NaOH, Sigma) (pore formation assays) using five freeze-and-thaw cycles and extruded 41x through a membrane with 400 nm pores. For the pore formation assays the LUVs were passed through a desalting column (Econo-Pac 10DG Desalting Columns, Biorad) equilibrated in outside buffer (20 mM HEPES, pH 7.0, 140 mM NaCl, 1 mM EDTA) to remove non-trapped calcein and diluted to a final maximum concentration of 1.25 mg/ml.

### Pore formation assays in LUVs

Pore formation assays were performed at 37°C in outside buffer with maximum 0.02 mg/ml LUVs and protein concentrations of 25 nM. The fluorescence was monitored over time (excitation: 495 nm, emission: 520 nm), observing the fluorescence increase of the calcein trapped in LUVs due to the dequenching after being released in the surrounding buffer upon pore formation. To determine the maximum fluorescence (100% value), triton-X-100 (max. 0.05%) was added at the end of the fluorescence kinetic measurement. In all protein combinations, the inhibitor (Bcl-xL) was added first, then the pore former (Bax) and finally, the activator (cBid) in a 1:1:1 stoichiometric ratio.

### Mitochondria purification and cytochrome c release assays

Crude mitochondria were isolated from mouse liver as described in literature[30]. In short, fresh mouse livers with a total of approximately 3.2 g were cut in small pieces on ice and homogenized in sucrose buffer using a glass dounce homogenizer. The homogenate was centrifuged at 600*×g* for 10 minutes at 4°C to remove any cell debris. Subsequentially, the supernatant was centrifuged at 7000*×g* for 10 minutes at 4°C. Approximately 1 mL of crude mitochondria suspension in sucrose buffer was then obtained and kept on ice. Mitochondrial proteins were considered as an index of mitochondrial concentration, and they were determined using a Bradford assay. The final protein concentration for mitochondria extraction amounted around 90 mg/mL.

For mitochondria extraction different mouse genotypes were used depending on the availability:

- IL-33 floxed, 1x CAS-9
- C57BL/6 SOPF
- Etv1-CreERT2
- SALSA(Stim1; Stim2 dKo) CEBP-Cre
- ERMP1

In order to test the direct and indirect inhibition of Bax activation by Bcl-xL, cytochrome c release assays were performed. Bax (7 *µ*M), cBid (0.85 *µ*M), and/or Bcl-xL (0.85 *µ*M) were mixed on ice with an aliquot of mitochondria suspension containing 22.64 mg/mL of proteins. To avoid mitochondrial disruption due to osmotic damage, the ratio between the protein buffer and the sucrose buffer was kept as 1:1. The samples were incubated at 37° C for 20 min and centrifuged for 3 minutes at 7000*×g* and 4° C. Then, 20 *µ*L of supernatant were diluted with 40 *µ*L of sucrose buffer and centrifuged for 15 minutes at 16,900*×g* and 4° C.

The absorbance spectrum of the supernatant was collected between 260 nm and 800 nm. In order to evaluate the absorbance of the Soret peak, the spectra were background corrected by an exponential fit (see fig. S3). The data processing was performed on MATLAB R2023b. For statistical analysis the maximum of the traces was used instead of a specific wavelength due to absorption shifts caused by the ratio of oxidized *vs.* reduced cytochrome c[31].

### Pellet and supernatant preparation

All pellet/supernatant samples analyzed (SDS PAGE, CW EPR) have been prepared as described in fig. 4A. Proteins and LUVs were mixed with a final protein concentration of 10-20 *µ*M and a final lipid concentration of 1-5 mg/ml. After one hour incubation at 37°C, the membrane and water sample were separated via centrifugation (120000xg). The pellet was resuspended and adjusted according to the methods requirements.

### Continuous wave EPR

All continuous wave (cw) EPR experiments were performed on a Bruker E500 X-band equipped with a super high Q cavity ER 4122 SHQ with a power of 2 mW and a modulation amplitude of 0.1 or 0.15 mT.

For the time-resolved measurements, four spectra were averaged before moving to the next time point. The temperature was controlled at 37°C with a nitrogen flow cryostat and a temperature controller (Eurotherm). Samples were measured in a total volume of 20 µL in glass capillaries (Blaubrand). Proteins at 10-20 *µ*M final concentrations were mixed on ice and introduced into the glass capillary. In a second step, a small volume of LUVs was added to the capillary. The first spectrum was detected after 2.5 min from the addition of LUVs to the protein mixture.

### Negative stain electron microscopy and data analysis

3 *µ*l of oligomeric Bcl-xL (0.05 mg/ml) were applied to a freshly glow discharged 400-mesh carbon-coated copper grid and incubated for 1 min before staining with 0.75% uranyl formate solution for 1 min. Grids were air dried for 5 min before they were imaged in a J1400 TEM (Jeol) equipped with a TVIPS 4k x 4k F416 detector.

36 micrographs recorded at 80,000x magnification (pixel size: 1.36 Å/px) were imported in CryoSPARC (version 4.7.0) and processed in negative stain mode with constant CTF. Particles were identified using the blob picker set to a diameter between 70 and 150 Å. 3,664 particles were extracted at a pixel size 2.72 Å/px and subjected to 2D classification using 36 classes to distinguish between intact oligomers and monomers.

## Author contributions

Christina Elsner: Conceptualization, Data curation, Formal analysis, Investigation, Methodology, Software, Validation, Visualization, Writing – original draft, Writing – review& editing. Ludovica M. Epasto: Formal analysis, Data curation, Writing – review & editing. Adeline Cieren: Methodology. Dominik Gendreizig: Methodology, Writing – review& editing. Svetlana Kucher: Methodology, Supervision, Writing – review& editing. Daniel Roderer: Formal analysis, Data curation, Writing – review & editing. Enrica Bordignon: Conceptualization, Supervision, Validation, Funding acquisition, Writing – original draft, Writing – review & editing.

## Data availability

All primary data are available upon request.

## Acknowledgements

Technical support for protein purifications was provided by O. Vadas, R. Visentin, L. Falconet (Protein Platform, Faculty of Medicine, University of Geneva, Switzerland). Technical support for mass photometry was provided by Y. Duhoo (Protein Production and Structure Core Facility, EPFL, Lausanne, Switzerland). Technical support during preliminary work was provided by J. Beermann and S. Bleicken (Ruhr University Bochum, Germany). EB and CE want to thank Prof. Dr. A. J. García-Sáez for providing the Bcl-2 plasmids. This work is supported by the SNSF Sinergia grant CRSII5_216587.

## Supplementary Information

**Figure S1:**
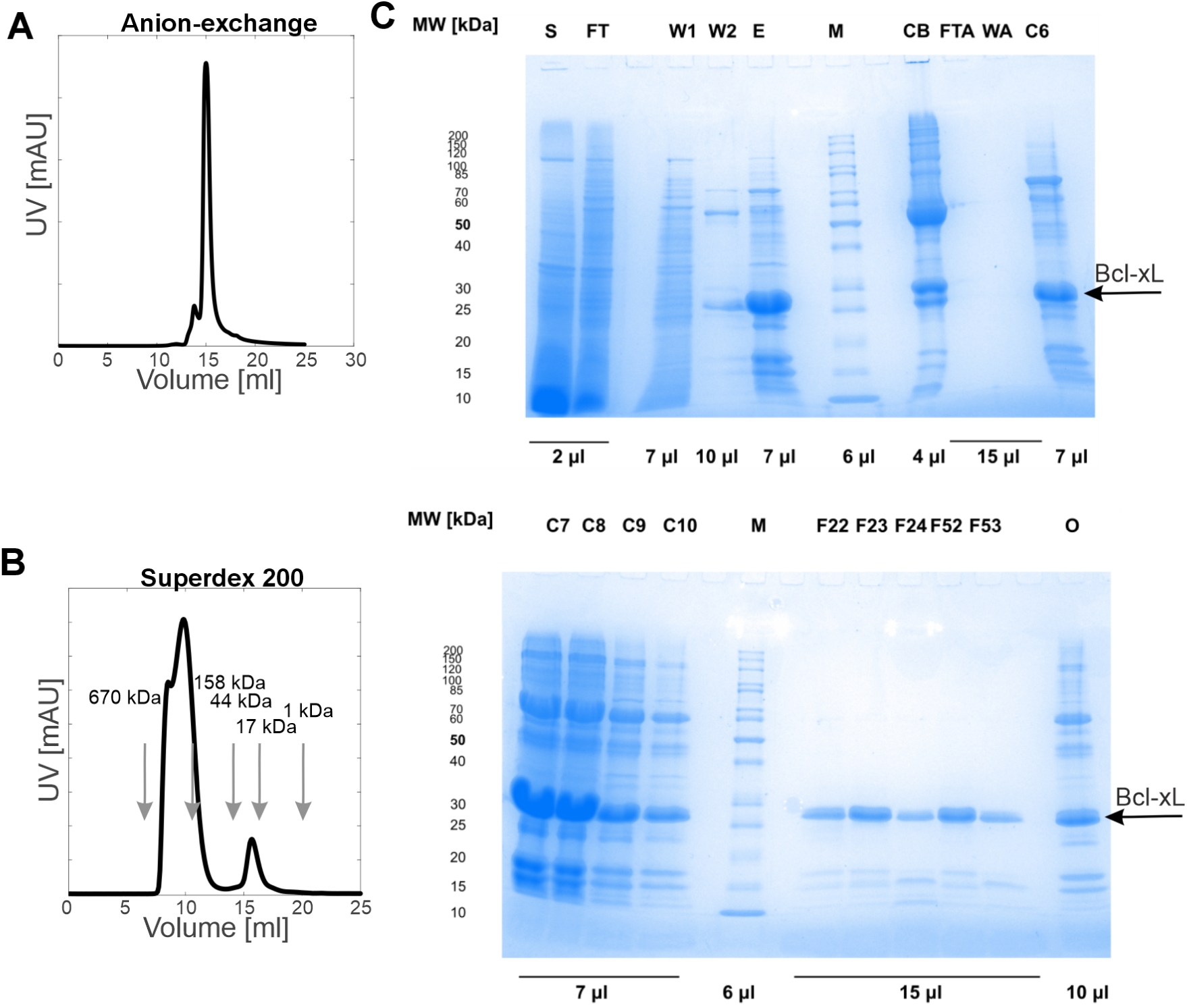
Bcl-xL 3-step purification. **A:** Anion-exchange chromatography detected at 280 nm during the Bcl-xL WT purification. The highest peak was pooled (2 ml total, fractions B6-B3) and used for further steps. SDS PAGE samples were prepared from all pooled fraction and the two neighboring fractions (B7 and B2) **B:** SEC analysis of the main peak eluted from the anion-exchange during Bcl-xL purification using a Superdex 200 10/300. Absorption detected at 280 nm. Since the maximum sample volume was 1 ml, this was run twice and the peaks considered together. The oligomer peaks were flash frozen and stored at −80°C, the monomer fractions were pooled, concentrated, aliquoted, and flash frozen before storage at −80°C. **C, D:** SDS PAGE analysis of the purification. S: Supernatant after cell-lysis. FT: flow through of the cell lysate loading onto the chitin beads. W1, W2: washing steps of the chitin beads. E: Elution of Bcl-xL WT off of the chitin beads after tag cleavage. M: Ladder. CB: Analysis of the chitin beads after elution but before cleaning. FTA: Flow through of the anion-exchange column. WTA: Wash of the anion exchange column. C6-C10: Fractions collected during the anion exchange. F22-24 + F52-F53: Monomer fractions after SEC and before pooling and concentrating. O: Pooled oligomer fractions. SDS PAGE samples were mixed with Laemmli buffer and heated to 95°C for 5 minutes. No reducing agents were used. Details of the purification are described in materials & methods.

**Figure S2:**
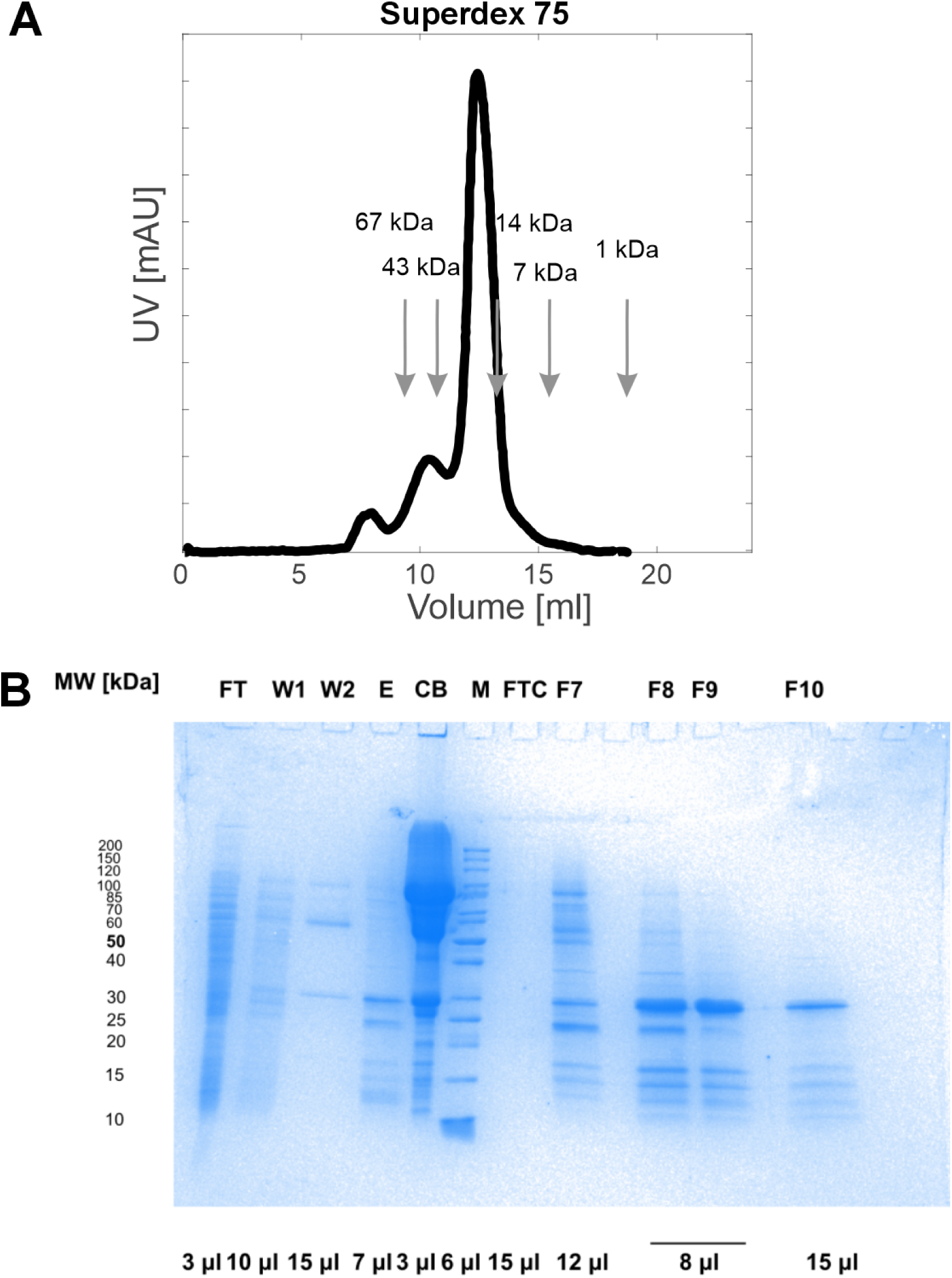
Bcl-xL 2-step purification (without anion exchange column). **A:** SEC of the concentrated chitin bead eluate on a Superdex 75 10/300 with indicated calibration. Absorption detected at 280 nm. The main peak was pooled, aliquoted, and flash frozen before storing at −80°C. SDS PAGE samples were taken from the main peak and the two neighboring fractions. **B:** SDS PAGE analysis of the purification. FT: flow through of the cell lysate loading onto the chitin beads. W1 + W2: washing steps of the chitin beads. E: Elution of Bcl-xL WT off of the chitin beads after tag cleavage. The 45 ml of eluate were concentrated to 1 ml for SEC analysis. CB: Analysis of the chitin beads after elution but before cleaning. M: Ladder. F7-F10: Fractions collected of the SEC. SDS PAGE samples were mixed with Laemmli buffer and heated to 95°C for 5 minutes. No reducing agents were used.

**Figure S3:**
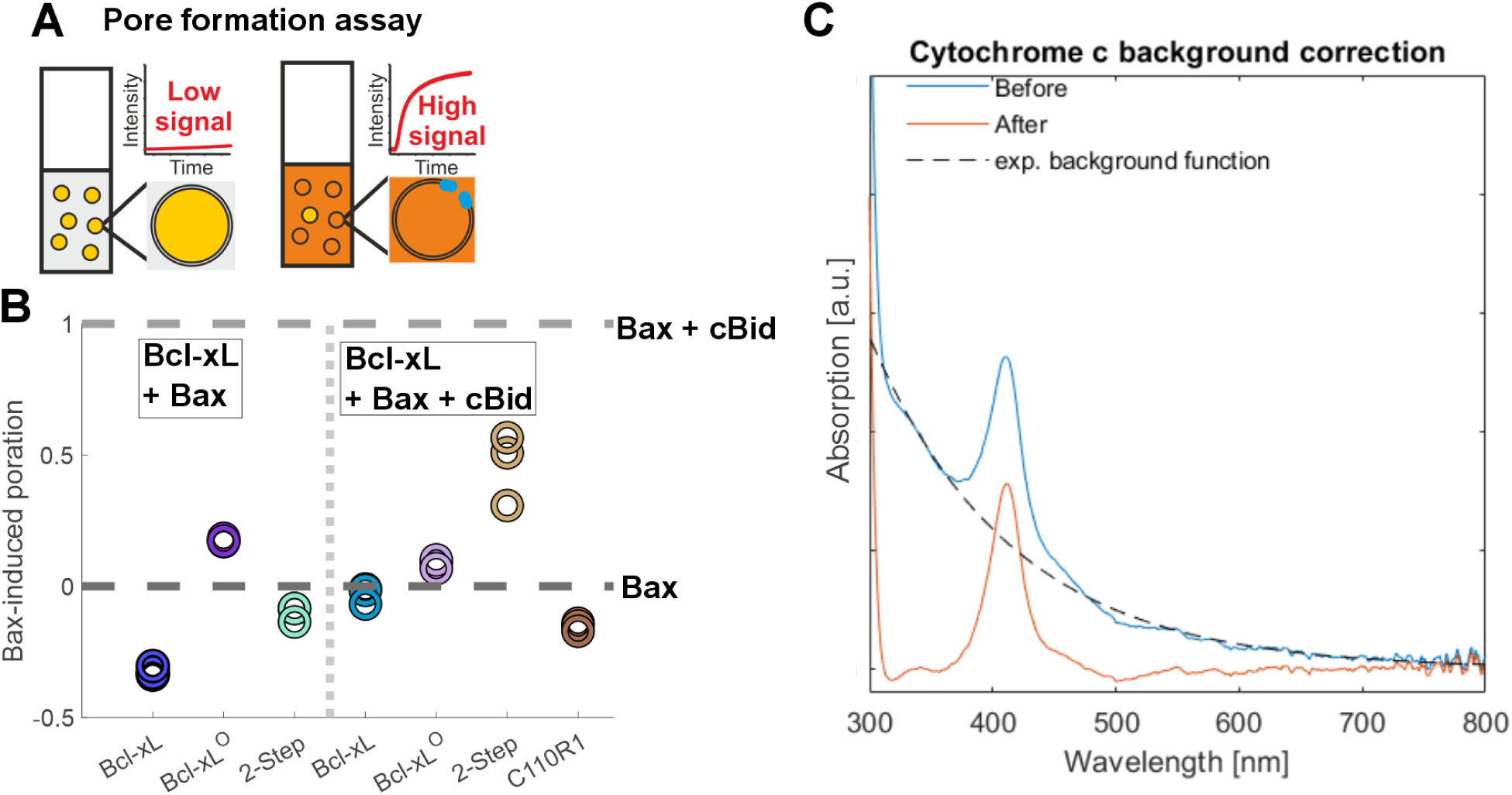
Activity assays. **A:** Graphic explanation of *in vitro* pore formation assays. Adapted from[32]. When the calcein is trapped at high concentration (80 mM) in the LUVs, it self-quenches leading to a low fluorescence signal. Upon membrane permeabilization by Bax, calcein is diluted in the surrounding buffer, leading to an increase in fluorescence. **B:** Comparison of technical replicates of the inhibitory effects of different Bcl-xL variants after 30 min of incubation: 1 is the normalized maximal Bax activity in presence of cBid, 0 is Bax autoactivity. Bcl-xL and Bcl-xL^O^ are monomers and oligomers (see fig. 2) as prepared in the 3-step purification described in materials & methods and in fig. S1. Following the same protocol, also the Bcl-xLC110R1 labeled monomer variant was prepared. The sample named ‘2-Step’ is the monomer prepared in a 2-step purification as shown in fig. S2 showing a decreased inhibition of the Bax/cBid mixture as compared to the other monomers prepared with a 3-step purification. **C:** Example of background correction of the cytochrome c UVvis spectrum with a monoexponential background function.

**Figure S4:**
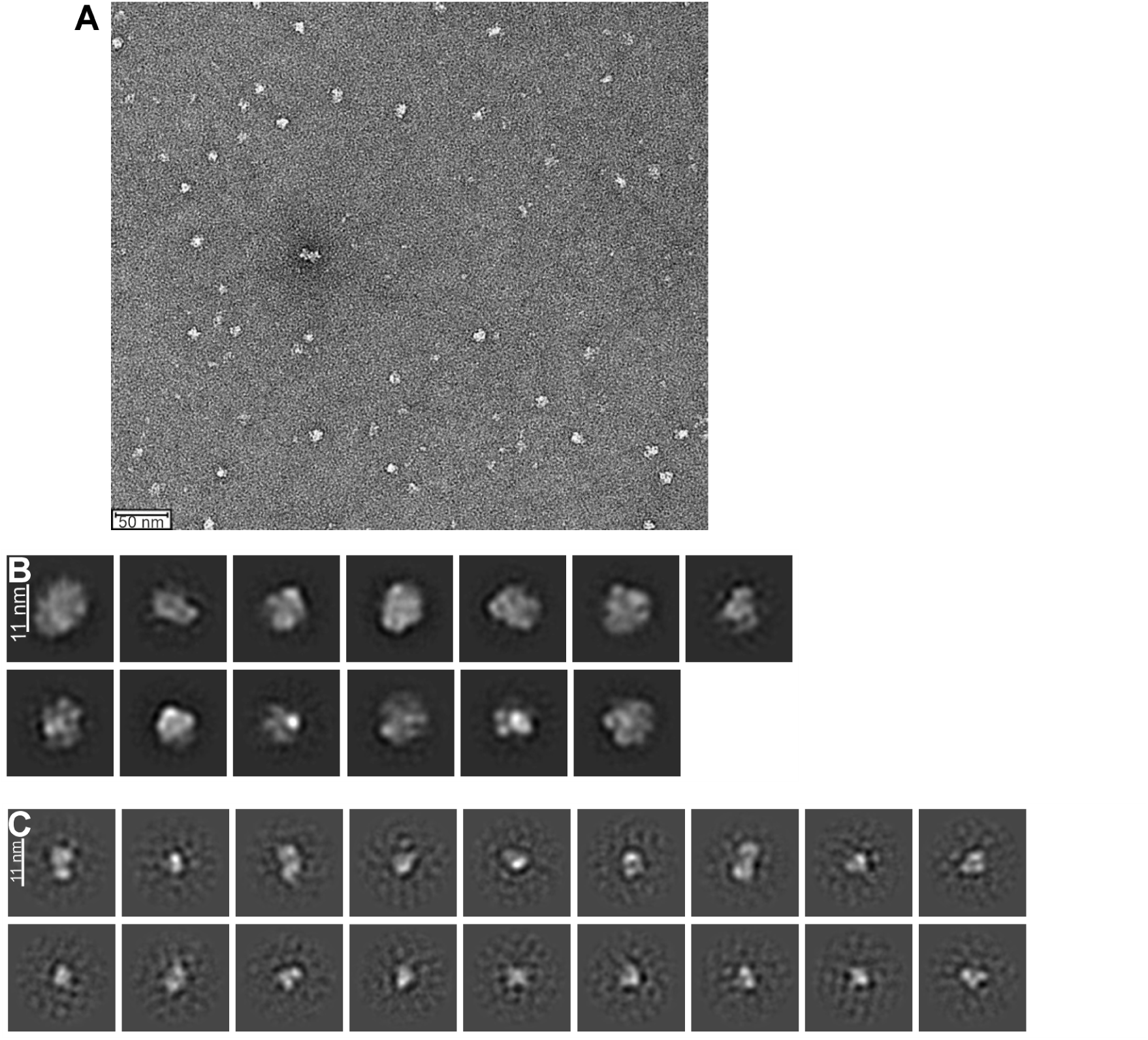
A: Representative EM micrograph of negatively stained of Bcl-xL^O^. B: 2D class averages of 1712 particles of negative stain data obtaine^d^ on Bcl-xL^O^. The size and shape of the observed particles are compatible with tetramers. C: 2D class averages of 1380 particles of negative stain data representing smaller particles, likely dimers and monomers.

**Figure S5:**
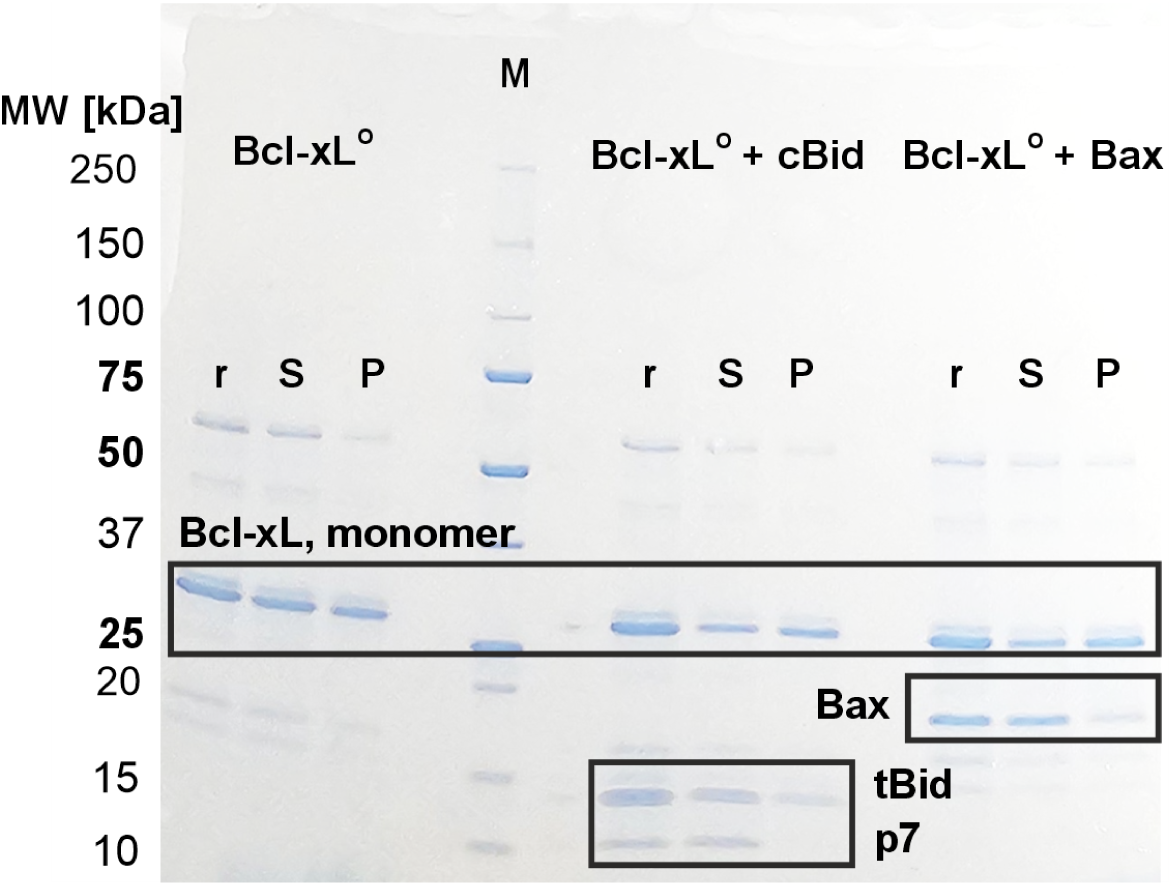
Analysis of water-membrane partitioning of oligomeric Bcl-xL. SDS PAGE analysis of Bcl-xL^O^ alone or in presence of cBid or Bax and LUVs. *P*: pellet obtained by the mixture with LUVs; *S*: supernatant obtained by the mixture with LUVs; *r*: reference protein mixture in solution without LUVs. Boxes highlight bands corresponding to monomeric Bcl-xL (26 kDa, upper panels), Bax (21 kDa, middle panel), cBid (tBid and p7 bands at 15 and 7 kDa, respectively, bottom panels). SDS PAGE samples were mixed with Laemmli buffer and DTT and heated to 95°C for 5 minutes.

**Figure S6:**
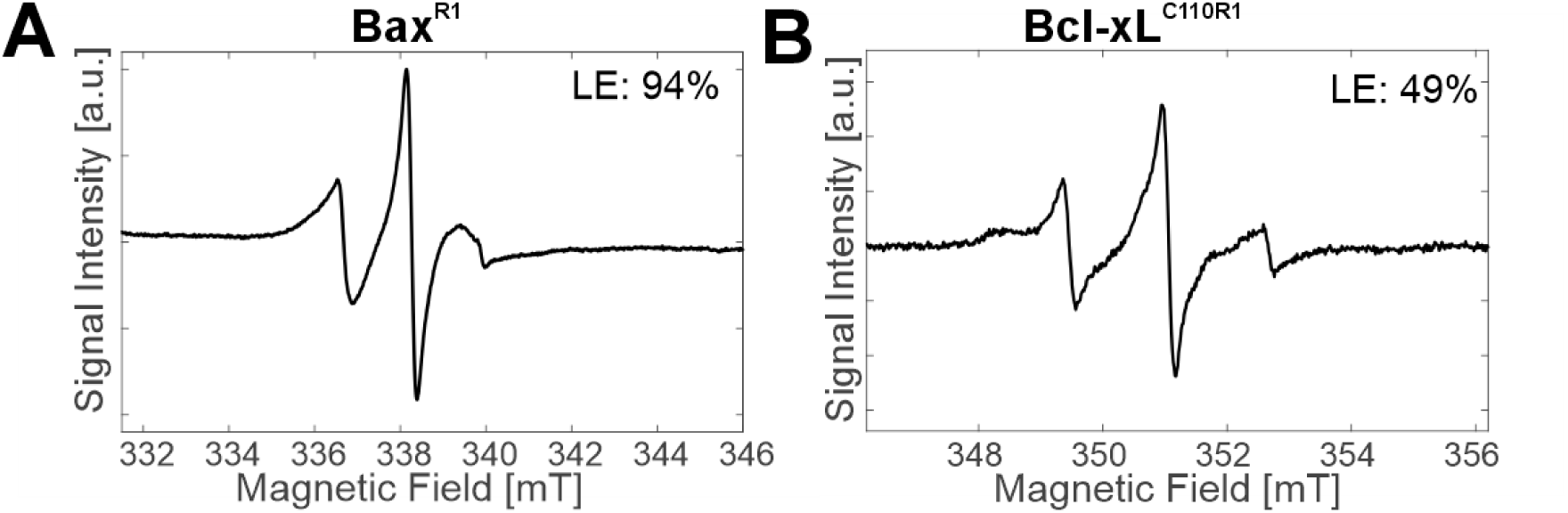
X-band cw EPR spectra. Measurements were performed at room temperature, labeling efficiencies (LE, calculated as spin per cysteine) of Bax^R1^ (**A**) and Bcl-xL^C110R1^(**B**) were calculated using the double integral of the spectra and the protein concentration obtained by UVvis.

## Bibliography

(1) Czabotar, P. E.; Lessene, G.; Strasser, A.; Adams, J. M. Nature reviews. Molecular cell biology 2014, 15, 49–63.

(2) Kale, J.; Osterlund, E. J.; Andrews, D. W. Cell death and differentiation 2018, 25, 65–80.

(3) Czabotar, P. E.; Garcia-Saez, A. J. Nature Reviews Molecular Cell Biology 2023, DOI: 10.1038/s41580-023-00629-4.

(4) Nguyen, D.; Osterlund, E.; Kale, J.; Andrews, D. W. Biochemical Journal 2024, 481, 903–922.

(5) Billen, L. P.; Shamas-Din, A.; Andrews, D. W. Oncogene 2008, 27 Suppl 1, S93–104.

(6) Westphal, D.; Kluck, R. M.; Dewson, G. Cell Death & Differentiation 2014, 21, 196– 205.

(7) Ding, J.; Mooers, B. H.; Zhang, Z.; Kale, J.; Falcone, D.; McNichol, J.; Huang, B.; Zhang, X. C.; Xing, C.; Andrews, D. W.; Lin, J. Journal of Biological Chemistry 2014, 289, 11873–11896.

(8) O’Neill, J. W.; Manion, M. K.; Maguire, B.; Hockenbery, D. M. Journal of Molecular Biology 2006, 356, 367–381.

(9) Czabotar, P. E.; Westphal, D.; Dewson, G.; Ma, S.; Hockings, C.; Fairlie, W. D.; Lee, E. F.; Yao, S.; Robin, A. Y.; Smith, B. J.; Huang, D. C. S.; Kluck, R. M.; Adams, J. M.; Colman, P. M. Cell 2013, 152, 519–531.

(10) Llambi, F.; Moldoveanu, T.; Tait, S. W. G.; Bouchier-Hayes, L.; Temirov, J.; McCormick, L. L.; Dillon, C. P.; Green, D. R. Molecular cell 2011, 44, 517–531.

(11) Kalkavan, H.; Green, D. R. Cell Death & Differentiation 2018, 25, 46–55.

(12) Thuduppathy, G. R.; Craig, J. W.; Kholodenko, V.; Schon, A.; Hill, R. B. Journal of Molecular Biology 2006, 359, 1045–1058.

(13) Bleicken, S.; Wagner, C.; García-Sáez, A. J. Biophysical journal 2013, 104, 421–431.

(14) Bleicken, S.; Hantusch, A.; Das, K. K.; Frickey, T.; Garcia-Saez, A. J. Nature communications 2017, 8, 73.

(15) Vargas-Uribe, M.; Rodnin, M. V.; Ladokhin, A. S. Biochemistry 2013, 52, 7901–7909.

(16) Vasquez-Montes, V.; Vargas-Uribe, M.; Pandey, N. K.; Rodnin, M. V.; Langen, R.; Ladokhin, A. S. Biochimica et Biophysica Acta (BBA) - Proteins and Proteomics 2019, 1867, 691–700.

(17) Kale, J.; Chi, X.; Leber, B.; Andrews, D. In Methods in Enzymology; Elsevier: 2014; Vol. 544, pp 1–23.

(18) Bogner, C.; Kale, J.; Pogmore, J.; Chi, X.; Shamas-Din, A.; Fradin, C.; Leber, B.; Andrews, D. W. Molecular Cell 2020, 77, 901–912.e9.

(19) Raltchev, K.; Pipercevic, J.; Hagn, F. Chemistry – A European Journal 2018, 24, 5493–5499.

(20) Ryzhov, P.; Tian, Y.; Yao, Y.; Bobkov, A. A.; Im, W.; Marassi, F. M. Biophysical Journal 2020, 119, 1324–1334.

(21) Aranovich, A.; Liu, Q.; Collins, T.; Geng, F.; Dixit, S.; Leber, B.; Andrews, D. W. Molecular cell 2012, 45, 754–763.

(22) Hsu, Y.-T.; Wolter, K. G.; Youle, R. J. Proceedings of the National Academy of Sciences 1997, 94, 3668–3672.

(23) Rajan, S.; Choi, M.; Nguyen, Q. T.; Ye, H.; Liu, W.; Toh, H. T.; Kang, C.; Kamariah, N.; Li, C.; Huang, H.; White, C.; Baek, K.; Grüber, G.; Yoon, H. S. Scientific Reports 2015, 5, 10609.

(24) Denisov, A. Y.; Sprules, T.; Fraser, J.; Kozlov, G.; Gehring, K. Biochemistry 2007, 46, 734–740.

(25) Bleicken, S.; Classen, M.; Padmavathi, P. V. L.; Ishikawa, T.; Zeth, K.; Steinhoff, H.-J.; Bordignon, E. The Journal of biological chemistry 2010, 285, 6636–6647.

(26) Bleicken, S.; Assafa, T. E.; Stegmueller, C.; Wittig, A.; Garcia-Saez, A. J.; Bordignon, E. Cell death and differentiation 2018, 25, 1717–1731.

(27) Flores-Romero, H.; Hohorst, L.; John, M.; Albert, M.-C.; King, L. E.; Beckmann, L.; Szabo, T.; Hertlein, V.; Luo, X.; Villunger, A.; Frenzel, L. P.; Kashkar, H.; Garcia-Saez, A. J. The EMBO Journal 2022, 41, e108690.

(28) Desagher, S.; Osen-Sand, A.; Nichols, A.; Eskes, R.; Montessuit, S.; Lauper, S.; Maundrell, K.; Antonsson, B.; Martinou, J.-C. The Journal of Cell Biology 1999, 144, 891– 901.

(29) Suzuki, M.; Youle, R. J.; Tjandra, N. Cell 2000, 103, 645–654.

(30) Frezza, C.; Cipolat, S.; Scorrano, L. Nature Protocols 2007, 2, 287–295.

(31) Schweitzer-Stenner, R. New Journal of Science 2014, 2014, 1–28.

(32) Teucher, M.; Zhang, H.; Bader, V.; Winklhofer, K. F.; García-Sáez, A. J.; Rajca, A.; Bleicken, S.; Bordignon, E. Scientific Reports 2019, 9, 3667.

